# Examining Horizontal Transmission of *Nippostrongylus brasiliensis* in Mice to Assess Biosecurity Risks

**DOI:** 10.1101/2023.01.28.526042

**Authors:** Rebecca J Floyd, Rodolfo J Ricart Arbona, Sebastian E Carrasco, Neil S Lipman

## Abstract

*Nippostrongylus brasiliensis* (Nb) infected mice are commonly used to study immune responses. There is no consensus regarding the biosecurity measures that should be employed when housing Nb-infected mice and rats. Transmission is reported to not occur when infected mice are co-housed with naive mice. We sought to determine what conditions, if any, favor horizontal transmission. Female NOD.Cg-*Prkdc*^*scid*^ *Il2rg*^*tm1Wjl*^/Sz (NSG; n=12) and C57BL/6J (B6; n=12) mice inoculated with 750 Nb L_3_ larvae were cohoused with naïve NSG (n=24) and B6 (n=24) mice at a ratio of 1 infected:2 naïve mice per cage for 28 days within static microsiolator (MI) cages changed on a 14-day interval. We also assessed whether larval development to the L_3_ stage could occur when Nb egg-containing fecal pellets are maintained under 4 different environmental conditions (dry, moist, soiled bedding, and control) and whether infection results when naïve NSG mice (n=9) are housed in MI cages with infective L_3_ larvae-spiked (10,000/cage) soiled bedding. NSG mice (n=3) were also gavaged with Nb eggs to model the potential for infection to occur following coprophagy. Both naïve NSG (9 of 24) and B6 (10 of 24) mice cohoused with an infected cagemate passed Nb eggs in feces as early as 1 day and intermittently for varying periods following cohousing, presumably a result of coprophagy as adult worms were not detected at euthanasia. While eggs were able to develop into L_3_ larvae under 2 environmental conditions (moist and control), none of the NSG mice housed in cages with L_3_-spiked bedding or gavaged with eggs became infected with Nb. Findings demonstrate that horizontal transmission does not occur when mice are housed with Nb-shedding cagemates in static MI cages with a 14-day cage changing interval. Results from this study can be used to inform biosecurity practices when working with the Nb-infected mouse model.

## INTRODUCTION

The mouse model of *Nippostrongylus brasiliensis* (Nb) infection is a well-established model for studying systemic and mucosal immune responses.^3; 5; 6; 18; 25; 38^ An extensive list of publications, dating back to the early 1900’ s, describe the parasite’ s life cycle, biology, and immunophysiology.^3; 5; 6; 18; 25^ Research utilizing the Nb mouse model has been instrumental in understanding the molecular mechanisms of Th2 type immune responses as well as the role it plays in protective immunity against helminth infections.^2; 5; 6; 16;^ Nb is commonly propagated by passaging through rats because large quantities of eggs and L_3_ larvae can easily be collected from fecal pellets or by removal of cecal and colonic contents.^6; 25^ Alternatively, Nb can be propagated through mice with the Swiss Webster stock most commonly used.^6^ Mice, rather than rats, are often studied due to the extensive characterization of inbred and the availability of genetically engineered strains.^25^ The most commonly used mouse strains are the C57BL/6 and BALB/c.^6^ Subcutaneous inoculation with 500 (mouse-adapted) or 750 (rat-adapted) L_3_ larvae is recommended when using the Nb mouse model.^6^

The life cycle of Nb is direct and approximately 2 weeks in length.^3; 5; 6; 18; 25^ Eggs are ellipsoidal with a thin shell measuring 50 to 70 by 27 to 40 µm.^6; 25^ They are excreted in the feces as 16 to 20 cell embryos.^6^ First stage larvae (L_1_) hatch within 18 to 24 hours at 18 to 26°C.^6; 25^ Over the following 5 to 6 days, larvae undergo 2 additional molts before forming unsheathed, filariform, infective third stage (L_3_) larvae measuring 620 to 750 µm in length. Infection occurs through direct skin penetration or by ingestion of L_3_ larvae. Skin penetration occurs within 5 minutes of contact, larvae then migrate through the epidermis into blood vessels to reach the lungs by 11 hours post-infection (PI). The 3^rd^ molt occurs in the lungs between 19 to 32 hours PI forming the 4^th^ stage (L_4_) larvae. L_4_ larvae remain in the lungs for up to 50 hours before migrating up the trachea, being coughed up and swallowed down the esophagus through the stomach reaching the small intestine where the final molt occurs to form L_5_ immature adults, 90 to 108 hours PI.^2; 3; 5; 6; 18; 25^ There, adult nematodes become sexually mature and reach their final size (3 to 4.5 mm for males and 4 to 6 mm for females). Gravid females can first be detected at the end of the 5^th^ day and eggs can be found in the feces approximately 5 to 6 days PI. In immunocompetent mice and rats, maximum egg production occurs 6 to 9 days PI followed by a gradual decline and worm expulsion on day 10 to 15.^2; 3; 5; 6; 18; 25^

Once Nb infection is cleared, lifelong immunity persists in immunocompetent animals.^5; 6; 19^ However, there are strain differences in susceptibility to infection. Th2-skewed strains, such as the BALB/c, are more resistant as compared to C57BL/6, a Th1-skewed strain that shed higher numbers of eggs.^43^ However, both strains expel the parasite by day 13 PI.^43^ In contrast, immunodeficient mice, e.g., athymic nude, that can’ t mount a T-cell response may fail to expel nematodes completely and become chronically infected.^1; 14; 21; 22; 31^

Infection in humans is reported to be self-limiting following skin penetration by L_3_ larvae, therefore donning proper PPE, including a laboratory coat and gloves, is recommended when working with Nb.^6^ Consideration must also be given to horizontal transmission to susceptible research animals when using this model. This risk is greatest when infecting immunocompromised mice that become chronically infected and shed high numbers of eggs. Biosecurity measures vary by institution, with infected rodents maintained following ABSL-1 or ABSL-2 practices, as there is not a definitive published study examining horizontal transmission when housing Nb-infected mice and rats. Recommendations for animal facility containment are described by *Camberis et al*. (2003) which states horizontal transmission should not occur in mice, even when infected and naïve animals are co-housed, if the cage is changed twice weekly as the infective L_3_ larvae stage require 5 to 6 days under optimal conditions of temperature and humidity to develop.^6^ While twice weekly cage changing is recommended, even longer periods are reported to be acceptable due to desiccation of larvae by low humidity, HEPA-filtered air at the cage level.^6^ Cage cleaning is also reported to prevent transmission as eggs and larvae are susceptible to hot water and detergent.^6^ However, references are not provided for these statements and presumably reference unpublished data.

The purpose of this study was to determine, using multiple strategies and ideal environmental conditions (for Nb development), whether horizontal transmission can occur in Nb-infected mice when cohoused with naïve mice, and whether housing conditions, i.e., cage change frequency, mouse strain, and immune status play a role in transmission. In addition, we sought to determine if L_3_ larvae can develop when fecal pellets containing Nb eggs are maintained under different environmental conditions and whether transmission occurs when naïve immunocompromised mice are housed in cages spiked with a large number of infective L_3_ larvae. Findings from this study can be used by institutions to determine the appropriate biosecurity measures to be implemented when using this model.

## MATERIALS AND METHODS

### Study Design

Three types of experiments were undertaken to evaluate the likelihood of horizontal Nb transmission.

#### Intracage Transmission

C57BL/6J (B6; n=12) and NOD.Cg-*Prkdc*^*scid*^ *Il2rg*^*tm1Wjl*^/SzJ (NSG; n=12) mice were inoculated with 750 L_3_ larvae. Egg shedding was evaluated daily in inoculated mice beginning on day 5 through day 35 days post inoculation (DPI), except for B6 mice in which daily egg counts were reduced to daily determination in 6 mice beginning on day 7 DPI. A greater number of NSG mice were evaluated as there is no published historical data regarding egg shedding patterns in this strain, whereas egg shedding in B6 mice is well documented.^6; 34; 43^ On day 7 DPI (B6 peak egg shedding), the Nb-inoculated and confirmed shedding B6 (n=12) and NSG (n=12) mice were randomized and cohoused with naïve B6 (n=24) or NSG (n=24) mice at a ratio of 1:2 (infected: naïve) mice per cage as detailed in Table 1. Each cohort of 4 groups was repeated 6 times yielding a total of 6 Nb-inoculated and 12 naive cagemates per group. Eggs per gram (EPG) of feces were determined for the naïve B6 and NSG mice starting at day 7 until day 28 of cohousing for the first 2 repetitions. After eggs were observed unexpectedly at 7 days post cohousing (DPH), coprophagy was suspected and egg counts were performed starting at 1 DPH for the following 4 repetitions (n=32) to ascertain whether eggs were present at earlier timepoints. Nb-inoculated mice were euthanized via carbon dioxide on 35 DPI and naïve mice at 28 DPH, and serum was collected for IgE determination (inoculated and naïve B6 mice only; n=24); gastrointestinal tract, lungs, heart, spleen, and lymph nodes were removed for histopathology; and, adult L_5_ worm burdens were quantified.

**Table 1:** Cohousing Configurations

| Group | Inoculated | Naïve |
| --- | --- | --- |
| 1 | 1 B6 | 2 B6 |
| 2 | 1 B6 | 2 NSG |
| 3 | 1 NSG | 2 B6 |
| 4 | 1 NSG | 2 NSG |

#### Cage Inoculation

To determine if mice can be infected with Nb when a high burden of infective L_3_ larvae are present in the bedding, naive NSG mice (n=9) were randomly selected and housed 3 per cage. Seven days later, each cage containing soiled bedding was spiked with 10,000 L_3_ larvae (∼1,000 larvae/ml) distributed throughout the cage by applying a 0.84 ml aliquot of PBS suspended larvae onto the top of the bedding 12 times, evenly spaced following a 3 × 4 pattern. Seven days after spiking the bedding with larvae, cages were changed, and 1-5 fecal pellets were collected daily directly from each mouse for 21 days. After no eggs were identified, cages were reinoculated a second time with 10,000 L_3_ larvae on day 21 as previously described and fecal pellets were collected for floatation daily until day 35, at which time the mice were euthanized, tissue samples collected for histopathology, and worm burdens quantified.

#### In vitro Larval Development

To determine if L_3_ larvae can develop when fecal pellets containing Nb eggs are incubated under environmental conditions replicating intracage conditions, approximately 1 gram of feces pooled from 3 Nb-shedding NSG mice were plated on 100 mm Petri plates (Falcon®, Corning, NY) containing either 5 grams of autoclaved aspen chip bedding (PWI Industries, Quebec, Canada; dry), 5 grams autoclaved aspen chip bedding mixed with 5 ml sterile water (moist), or 5 grams of previously autoclaved aspen chip bedding collected from a cage which housed 5 mice for 7 days (soiled; n=3 Petri plates per condition: dry, moist, and soiled). Control plates (n=3) contained autoclaved activated charcoal and approximately 1-gram Nb-containing fecal pellets softened in sterile water and plated on a piece of moistened 90 mm filter paper (Whatman™, Cytiva, Marlborough, MA) as described below. The control plates replicated optimal conditions under which the parasite is cultured *in vitro*. All 100 mm plates were placed into larger 150 mm Petri plates (Falcon®) containing enough sterile water to completely cover the bottom of plate and incubated at room temperature (∼26°C) for 7 days in a humidified (tray placed at the bottom of the incubator containing 200 ml of sterile water) incubator (Model No. 100A, Blue M, Blue Island, IL). Plates were removed from the incubator every 3 days and the lids removed for 1 minute to allow for aeration. Groups that were required to remain moist were checked every 3 days, and approximately 2 ml sterile water added to prevent plates from drying. On day 7, the contents of each 100 mm plate were evaluated for L_3_ larvae using the Baerman technique.

### Supplemental Studies Conducted to Investigate Unexpected Results

#### Nb Inoculation of Athymic Nude Stocks

Nb inoculated athymic nude mice (n=6; nude-1) were initially used as the immunocompromised mouse in the study as they were reported to chronically shed Nb eggs.^1; 21; 22; 31^ However, due to inconsistent infection, low fecal EPG and cessation of egg shedding at 10 DPI, they were replaced with NSGs as previously described and reflected in Table 1. An alternate athymic nude mouse stock (n=6; nude-2) from another vendor was inoculated with Nb L_3_ to determine if the results were attributable to genetic differences between stocks. All mice were inoculated with Nb L_3_, daily fecal samples collected and fecal floats performed until mice were euthanized at 35 DPI, at which point tissues were collected for histopathology and adult worm burdens determined.

#### Gavage of Feces Containing Nb Eggs

Naïve NSG and B6 mice, co-housed with Nb-infected NSG mice, were found to have eggs via fecal flotation as early as 1 DPH. To confirm that coprophagy was the source and if it occurred, could eggs embryonate and hatch within the gastrointestinal tract, 3 NSG mice were gavaged with a solution containing Nb eggs. The solution was prepared by mixing 4 to 5 fecal pellets from an Nb egg-shedding NSG mouse with 1 ml of 50% dextrose solution. The resulting slurry was centrifuged at 1300 rpm for 5 minutes and the supernatant removed. A 100 µl aliquot of the supernatant was used to confirm the presence of Nb eggs and enumerate the sample. The remainder of the sample (900 µl) was divided in thirds, each of which was diluted in sterile water (1:1) and administered to an NSG mouse via gavage. Mice were cohoused in a single wire bottom cage containing a moist paper towel overnight. Approximately 5 grams of fecal pellets were collected the following morning and examined for the presence of eggs via flotation. As the collected samples were initially Nb egg negative, the mice were gavaged a second time the following day with the same sample preparation. Following the second gavage, fecal pellets were collected daily for 11 days, and fecal flotation performed. On the final day, mice were euthanized, and a complete necropsy performed to identify the presence of adult worms.

### Experimental Animals

#### Mice

Six to 8-week-old female C57BL/6J (B6; The Jackson Laboratory [JAX], Bar Harbor ME), NOD.Cg-*Prkdc*^*scid*^ *Il2rg*^*tm1Wjl*^/SzJ (NSG; JAX), J:NU (nude-1; JAX), and Hsd:Athymic Nude-*Foxn1*^*nu*^ (nude-2; Envigo, Indianapolis, IN) were used. All mice were free of mouse hepatitis virus, Sendai virus, mouse parvovirus, minute virus of mice, murine norovirus, pneumonia virus of mice, Theiler meningoencephalitis virus, epizootic diarrhea of infant mice (mouse rotavirus), ectromelia virus, reovirus type 3, lymphocytic choriomeningitis virus, K virus, mouse adenovirus 1 and 2, polyoma virus, murine cytomegalovirus, mouse thymic virus, Hantaan virus, mouse kidney parvovirus, murine astrovirus-2, *Mycoplasma pulmonis, Citrobacter rodentium, Salmonella* spp., *Filobacterium rodentium, Clostridium piliforme, Corynebacterium bovis, Chlamydia muridarum*, fur mites (*Myobia musculi, Myocoptes musculinis*, and *Radfordia affinis*), pinworms (*Syphacia* spp. and *Aspiculuris* spp.), and *Encephalitozoon cuniculi* when the studies were initiated, as determined by testing naïve outbred Swiss Webster (Tac:SW) mice exposed repetitively to soiled bedding from cages housing mice in the colony.

#### Rat

An 8-week-old male Sprague Dawley (Crl:SD [SD]; Charles River Laboratories, Wilmington, MA) was used for generating L_3_ larvae for the studies. The rat was free of Sendai virus, Hantaan virus, Kilham rat virus, lymphocytic choriomeningitis virus, mouse adenovirus, pneumonia virus of mice, rat minute virus, rat parvovirus, reovirus type 3, Sendai virus, Theiler murine encephalomyelitis virus, Toolan H1 virus, *Filobacterium rodentium, Citrobacter rodentium, Clostridium piliforme, Corynebacterium kutscheri, Mycoplasma pulmonis, Pasteurella pneumotropica, Salmonella* spp., *Streptobacillus moniliformis, Encephalitozoon cuniculi*, fur mites, and pinworms (*Syphacia* spp. and *Aspiculuris* spp.) as determined by testing naïve outbred Sprague Dawley (Crl:SD) rats exposed repetitively to soiled bedding from cages housing rats in the colony.

### Husbandry and Housing

Mice were housed in polysulfone static microisolator cages with stainless-steel wirebar lids (Allentown LLC, Allentown, NJ). The rat was housed in an individually ventilated polysulfone microisolator cage (# 4, Thoren Caging Systems, Hazelton, PA). Animals were housed on autoclaved corncob bedding (Bed-o’ Cobs^®^, Andersons, Maumee, OH) and provided acidified (HCl) reverse osmosis purified water (pH 2.5– 2.8) in polyphenylsulfone bottles with stainless-steel caps and sipper tubes (Techniplast, West Chester, PA) and fed a closed-formula, γ-irradiated diet (LabDiet 5053, PMI, St Louis, MO) ad libitum. Each cage was provided with a Glatfelter paper bag containing 6 g of crinkled paper strips (EnviroPak, WF Fisher and Son, Branchburg, NJ) for enrichment. Rats were also provided a gnaw toy (Nylabone®, Neptune City, NJ; rat only). Cages were maintained under ABSL-2 conditions in a climate-controlled cubicle at 72 ± 2 °F (22.2 ± 0.5 °C), 30% to 70% relative humidity, and a 12:12 hour (0600:1800 h) light:dark cycle. Intracage temperature and relative humidity were also measured and recorded by placing a data logger (Edstrom Date Logger, Waterford, WI) between the wirebar lid and filter top in 1 cage from each of 3 randomly selected groups during 28 days of cohousing to confirm that environmental conditions at the cage level were optimal for development of L_3_ larvae. All animal use was approved by Weill Cornell Medicine’ s (WCM’ s) IACUC and maintained in accordance with *the Guide for the Care and Use of Laboratory Animals 8*^*th*^ *Edition*.^20^ All animal use was conducted in agreement with AALAS’ position statements on the Humane Care and Use of Laboratory Animals and Alleviating Pain and Distress in Laboratory Animals. WCM’ s animal care and use program is AAALAC-accredited.

### Generation of N. brasiliensis L_3_ Larvae

Nb L_3_ larvae (∼130,000) generated by serial passage through Sprague Dawley rats were kindly donated by the Rudensky Laboratory at the Memorial Sloan Kettering Cancer Center (MSK). Additional L_3_ larvae were generated by inoculating a Sprague Dawley rat with approximately 5,000 L_3_ in a 0.5 ml suspension of sterile PBS (Gibco™ PBS [10X], pH 7.4, Dublin, Ireland) subcutaneously into the intrascapular area using a 23-gauge needle (BD PrecisionGlide™ Single-use Needle, Franklin Lakes, NJ). At 6 DPI, the rat was placed in a barren cage lined with water-moistened paper towels (Wypalls®, Kimberly-Clark, Irving, Texas) containing a wire-grid floor.^6^ Water-moistened paper towels were replaced daily. The presence of Nb eggs in the feces was confirmed on 6 DPI. Approximately 50 fecal pellets were collected from 7 to 11 DPI, placed in a 50ml sterile conical tube (Falcon®, Corning, NY), and softened with sterile water.^6^ An equal volume of autoclaved activated granulated charcoal (Ebonex, Melvindale, MI) was added to form a coarse paste. The fecal and charcoal paste was evenly distributed over a moistened piece of 90 mm filter paper (Whatman™, Cytiva, Marlborough, MA) placed in a 100 mm Petri plate (Falcon®). Prepared Petri plates were placed into larger 150 mm Petri plates (Falcon®) filled with approximately 20 mls of sterile water, covered to provide elevated humidity, and were incubated at room temperature for approximately 7 days.^6^ Dishes were checked every 3 days to ensure they were sufficiently moist and aerated by removing the lid for 1-minute. Additional sterile water was added to maintain moisture as needed. L_3_ larvae were harvested beginning on day 7 by adding approximately 1 ml of water to the periphery of the smaller dish plate using sterile transfer pipette, gently swirling the dishes, followed by aspiration of the added water into a 50 ml conical tube (Falcon) as previously described.^6^ L_3_ larvae were washed by adding 20 mls of sterile water to the tube, the larvae allowed to settle, and the added water was gently aspirated. The larvae were washed 2 additional times with 40 mls of sterile water followed by 3 washes with 40 mls of sterile PBS (Gibco™ PBS [10X]). Larvae were counted under a dissecting microscope using a McMaster slide and stored in vented U-shaped 275 ml tissue culture flasks (Polystyrene tissue culture flasks with vented caps, Corning®, Corning, NY) at 2,000 L_3_/ml.

### Inoculation of B6 and NSG Mice with Nb Larvae

A 5 ml aliquot of L_3_ larvae collected from the tissue culture flask was placed in a 50 ml conical tube (Falcon) and allowed to settle to form a pellet. The supernatant was removed and 20 mls of sterile PBS added. The procedure was repeated 3 times and the final pellet resuspended in approximately 3 mls of sterile PBS.^6^ Larval counts were determined in a 100 μl aliquot using a McMaster counting slide. The larval concentration was adjusted using PBS to provide 3,750 L_3_ larvae/ml.^6^ The concentration determination was repeated 3 times to ensure accuracy. Mice were injected subcutaneously in the interscapular area with approximately 750 L_3_ suspended in 200μl sterile PBS (Gibco™ PBS [10X]) with a 23-gauge needle (BD PrecisionGlide™ Single-use Needle) as previously described.^6^

### Detection and Enumeration of Nb Eggs

Individual mice were lightly restrained over a disposable towel (Wypall®) and 1 to 5 fecal pellets collected. Pellets were placed in 5 ml microcentrifuge tubes (Eppendorf™, Hamburg, Germany) and their weight determined to calculate the eggs per gram of feces (EPG). They were then soaked in 2 mls of a sodium nitrate solution (Fecasol®, Vetoquinol, Fort Worth, Texas) for 15 to 20 minutes to soften followed by adding an additional 2 mls of the sodium nitrate solution. The contents of the tube were mixed again to form a homogenous slurry. An aliquot was immediately pipetted (Blood Bank Pipets, Copan Diagnostics, Murrieta, CA) into a McMaster counting chamber slide (FEC Source, Grand Ronde, OR) until the chamber was filled. After 5 minutes, the slide was examined under a compound microscope (Eclipse E200, Nikon, Feasterville, PA) at 10X for the presence of Nb eggs. Egg counts were performed 3 times with a different sample aliquot. The mean of the 3 counts was used to calculate the EPG.^6^ If eggs were not identified on any of the 3 slides examined, the sample was identified as negative. If eggs were only found outside of the counting chamber on any of the replicates, the sample was considered positive and recorded as less than 25 EPG, the sensitivity of the McMaster counting slide provided by the manufacturer.

### Collection and quantification of Adult Nb

Following euthanasia, the proximal half of the small intestine was excised and placed in a 100 mm Petri plate (Falcon®) containing 1 ml PBS. The intestine was sliced longitudinally revealing the mucosa. It was then cut into small approximately 3 to 5 mm lengths, placed in cheese cloth (Regency Wraps Natural Ultra Fine Cheesecloth™), submerged in a 50 ml conical tube (Falcon®) filled with 45 ml PBS, and incubated in a water bath for 2 hours at 37°C. Following incubation, tubes were observed for any adult worms that settled to the bottom and the small intestines examined for any worms left attached to the mucosa using a dissecting microscope. Worms were collected using a transfer pipette (Blood Bank Pipets, Copan Diagnostics, Murrieta, CA) and the total number collected were counted under a dissecting microscope using reported methods.^6; 25^

### Isolation and Counting of _L^3^_ Larvae Using the Baerman Technique

A Baerman apparatus, consisting of a funnel, plastic tubing, and a clamp was used. The contents from each Petri dish containing different environmental conditions were placed in cheese cloth (Regency Wraps Natural Ultra Fine Cheesecloth™, Dallas, Texas) cut into an 8 × 8 cm square which was tied with a piece of string to form a pouch attached to a wooden applicator. Warm PBS (37°C) was added to the Baerman apparatus until it reached approximately 1 cm below the rim of the funnel. Pouches were placed into funnels such that they were suspended by the wooden applicator and their contents were completely submerged. Additional PBS was added as needed until the pouches were completely submerged. After 24 hours, a 5 ml sample was collected into 15 ml conical tubes (Falcon®) from the Baerman apparatus by slowly releasing the clamp placed at the bottom of the tube. The samples were given 5 minutes to allow L_3_ to settle. The L_3_ larvae were aspirated and were counted under a dissecting microscope using a McMaster counting slide as previously desctibed.^6^

### Histopathology

After euthanasia by CO_2_ asphyxiation, the distal half of the small intestine (including jejunum and ileum), sections of the cecum and proximal, middle, and distal colon, mesenteric lymph nodes, all lung lobes, tracheobronchial lymph nodes, heart, and spleen were removed and fixed in 10% neutral buffered formalin. Tissues were processed in ethanol and xylene, and embedded in paraffin in a tissue processor (Leica ASP6025, Leica Biosystems). Paraffin blocks were sectioned at 5-μm thickness, stained with hematoxylin and eosin (H&E), and examined by a board-certified veterinary pathologist (SEC).

### Determination of Serum IgE Concentrations

Immediately following euthanasia, 0.5 to 0.7 mls of blood was collected via cardiocentesis or from the caudal vena cava from 8 Nb-inoculated B6 mice and 16 naïve B6 mice and placed in serum separator tubes (BD Microtainers, Becton Dickinson, Franklin Lakes, NJ). Clotted blood samples were centrifuged at 1500 rpm for 10 minutes and serum was collected and stored at -80°C degrees until analyzed. Total IgE concentrations were determined in previously frozen samples using a commercially available IgE ELISA kit (IgE Mouse uncoated ELISA Kit, Invitrogen-Thermo Fischer Scientific, Waltham, MA) following the manufacturer’ s instructions. In brief, microliter wells were coated with capture antibody (pre-titrated, purified anti-mouse antibody) in coating buffer, sealed, and incubated at 4°C overnight. The wells were aspirated and washed (1X PBS, 0.05% Tween™-20), blocked (20X PBS with 1% Tween™-20 and 10% BSA and deionized water in a 1:10 dilution), and incubated for 2 hours at 4°C. Wells were aspirated, washed, and serum samples prediluted to 1:50 and sample diluent were added in duplicate. Samples were then aspirated, and wells washed prior to the addition of pre-titrated, biotin-conjugated anti-mouse IgE monoclonal antibody. The plate was sealed with parafilm and incubated at room temperature on a microplate shaker at 400 rpm for 1 hour. Detection antibody was removed, the wells were washed, and pre-titrated streptavidin added. Detection enzyme was removed, and the wells were washed prior to addition of tetramethylbenzidine. The reaction was stopped by adding 1 M H_3_PO_4_ and plates were read at 450 nm (ELx808 Ultra from BIO-TEK Instruments, Inc. in Winooski, VT). The mean of duplicate samples of total IgE concentrations were determined in comparison to a standard curve.

### Statistical Analysis

Cumulative mean EPG shed by Nb-inoculated NSG and B6, and separately by nude-1 and nude-2 mice were compared using the Mann-Whitney U test with the Holm-Šídák method used to compare egg shedding on individual days. Total IgE serum concentrations in Nb-inoculated and naïve B6 mice were compared using a Mann-Whitney U test. One-way ANOVA was used to compare the cumulative mean EPG shed by naïve and Nb-inoculated mice and the 4 different environmental conditions under which eggs were incubated with the Tukey multiple comparison test performed to compare individual variables. *P* values of less than or equal to 0.05 were considered statistically significant. All statistical analyses were performed using Graph Pad Prism software (Graph Pad Prism version 9.4.1, La Jolla, CA).

## RESULTS

### Intracage Transmission

#### Egg Shedding Kinetics in Nb inoculated mice

All Nb-inoculated NSG and B6 mice achieved patency. The number of eggs per gram of feces (EPG) shed by B6 (n=6) and NSG (n=12) mice inoculated with Nb are shown in Figure 1. Eggs were first detected in the feces of most NSG (8 of 12; 66.7%) and B6 (10 of 12; 83.3%) on 6 DPI. Of the remaining mice, eggs were first observed on 7 DPI except for an NSG mouse in which eggs were first detected on 9 DPI. Cumulative mean EPG were significantly higher (*P* = 0.0001) in NSG mice as compared to B6 mice, 14,936+/-2686 vs 1,647+/-1223 (mean, +/-SEM) respectively. EPGs were higher in B6 mice compared to NSG mice on 7 and 8 DPI, but the differences were not statistically significant. EPG in B6 mice peaked at 7 DPI and no eggs were detected after 9 DPI. In contrast, peak shedding occurred at 9 DPI in NSG mice and, although EPG decreased over time, mice continued to shed eggs throughout the duration of the study.

**Figure 1.**
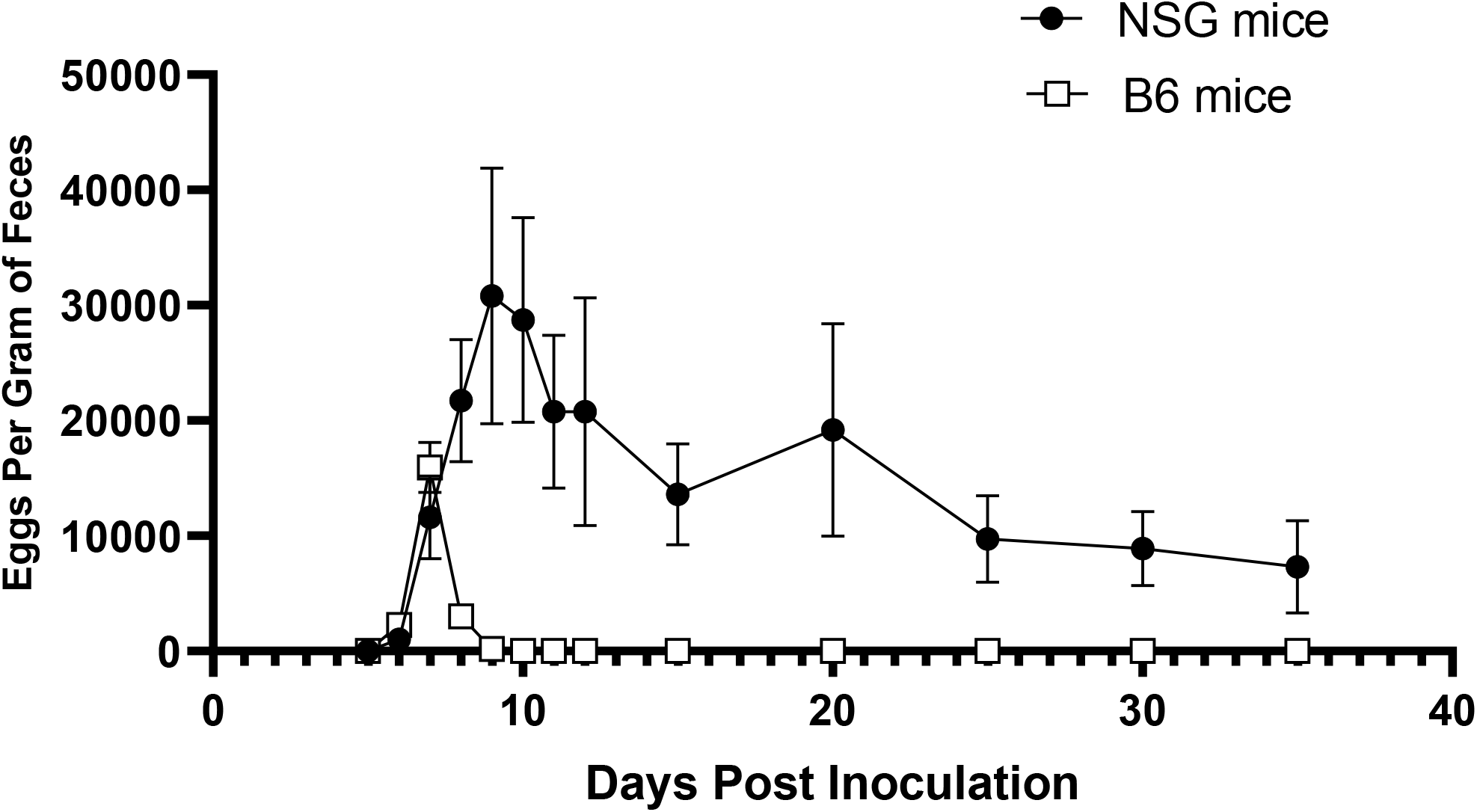
Data are shown as mean (+/-SEM) eggs per gram of feces for B6 (n=6) and NSG (n=12) mice inoculated with Nb on various days post inoculation (DPI) for the study duration of 35 DPI.

#### Horizontal transmission of Nb to Naïve B6 and NSG mice

Eggs were detected in varying numbers and intermittently in 10 of 12 (83 %) naïve B6 and 9 of 12 (75 %) naïve NSG mice cohoused with Nb-inoculated NSG mice during the 28-day evaluation period. Eggs were detected as early as 1 DPH in 2 of 12 B6 mice and 2 DPH in 2 of 12 B6 mice (Figure 2). The percentage of naïve B6 and NSG mice shedding eggs varied daily throughout the period of cohousing with Nb-inoculated NSG mice (Figure 3). The number of days during cohousing in which eggs were detected in the feces of individual naïve mice ranged from 0 to 17. No eggs were identified in fecal pellets of naïve NSG (n=12) or B6 (n=12) mice co-housed with Nb-inoculated B6 mice. Mean cumulative EPGs were higher in B6 mice as compared to NSG mice, but differences were not statistically significant. Naïve mice with the highest EPG were found to have been cohoused with Nb-inoculated NSG mice shedding the highest number of eggs. Neither lesions suggestive of Nb infection, nor larvae in the lungs, nor adult worms in the gastrointestinal tracts were identified in any naïve mice.

**Figure 2.**
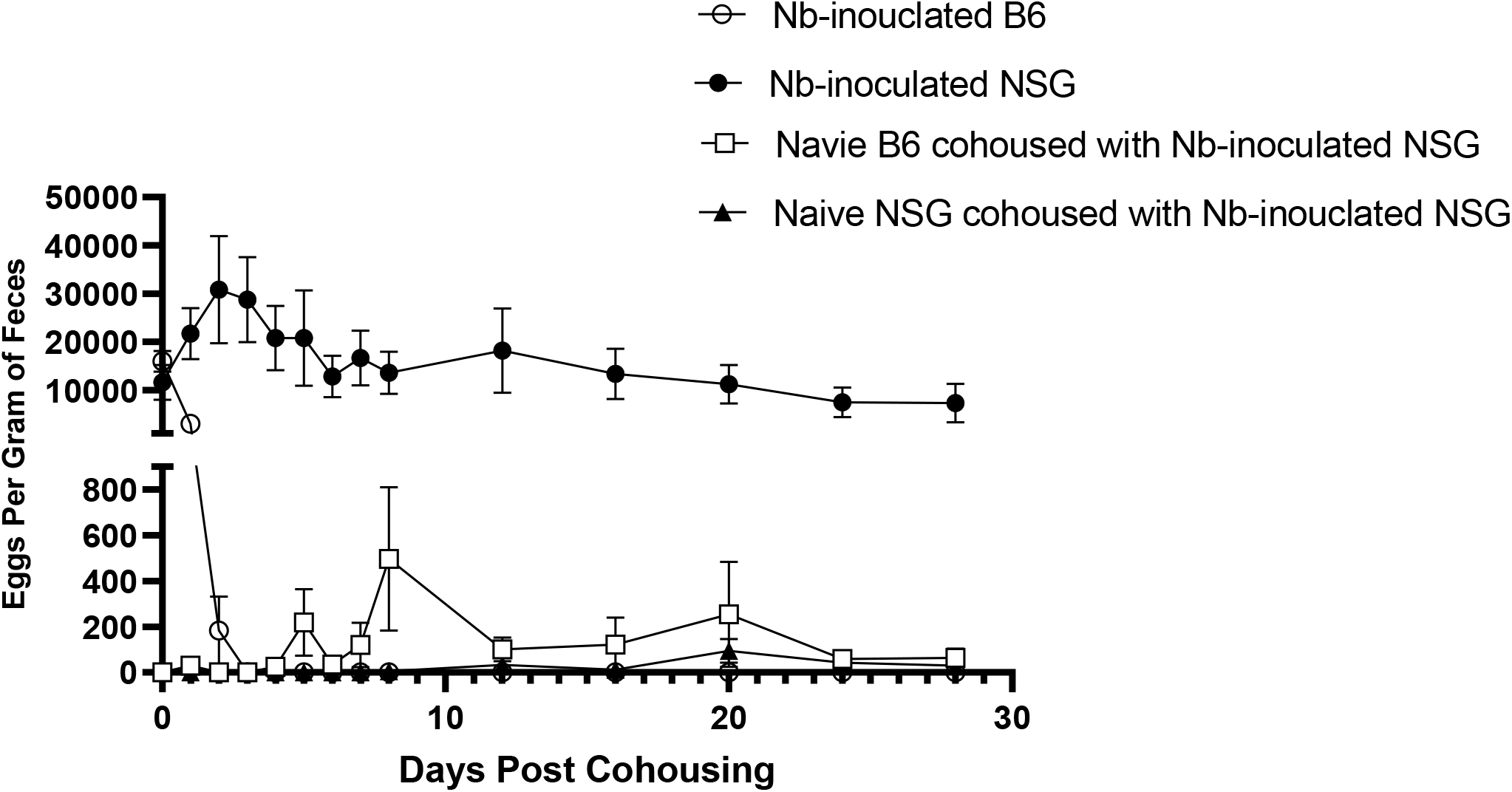
Data are shown as mean (+/-SEM) eggs per gram of feces for naïve B6 (n=12) and naïve NSG mice (n=12) cohoused with Nb-inoculated NSG (n=12) or Nb-inoculated B6 (n=6) mice on days 1 through 28 post-cohousing.

**Figure 3.**
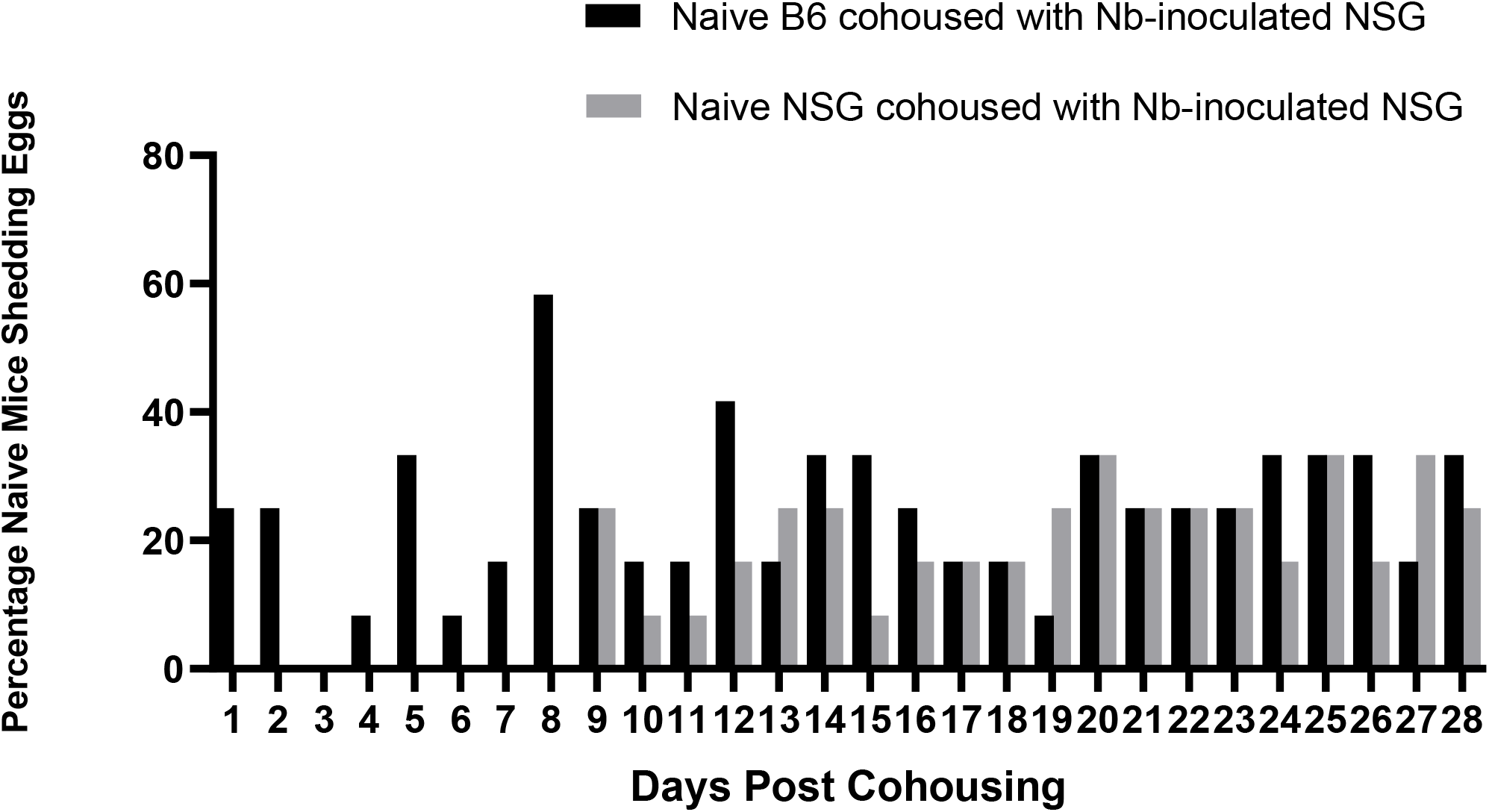
Data shown are the percentage of naïve NSG (n=8) and B6 (n=8) mice from 1 until 6 days post cohousing, and for all naïve NSG (n=12) and B6 (n=12) mice from 7 until 28 days post cohousing with eggs detected in feces while cohoused with Nb-inoculated NSG mice.

#### Adult (L_5_) worm counts

No adult worms were observed in Nb-inoculated B6 or naïve NSG and B6 mice. Adult worms (16.75+/-5.61; mean +/-SEM) were identified in the small intestines of all 12 Nb-inoculated NSG mice.

#### Serum IgE concentration

Median serum IgE levels were significantly elevated (*P*=<0.0001) on 28 DPH in Nb-inoculated B6 mice as compared to naïve B6 mice cohoused with Nb-inoculated NSG mice (Figure 4). Although a single naïve B6 mouse had an elevated IgE concentration (2,993.1 ng/ml) as compared to other naïve B6 mice (n= 15; 429.0 +/-89.8; mean +/-SEM), no eggs or adult worms were detected, nor were lesions suggestive of an Nb infection seen on histopathology in this mouse.

**Figure 4.**
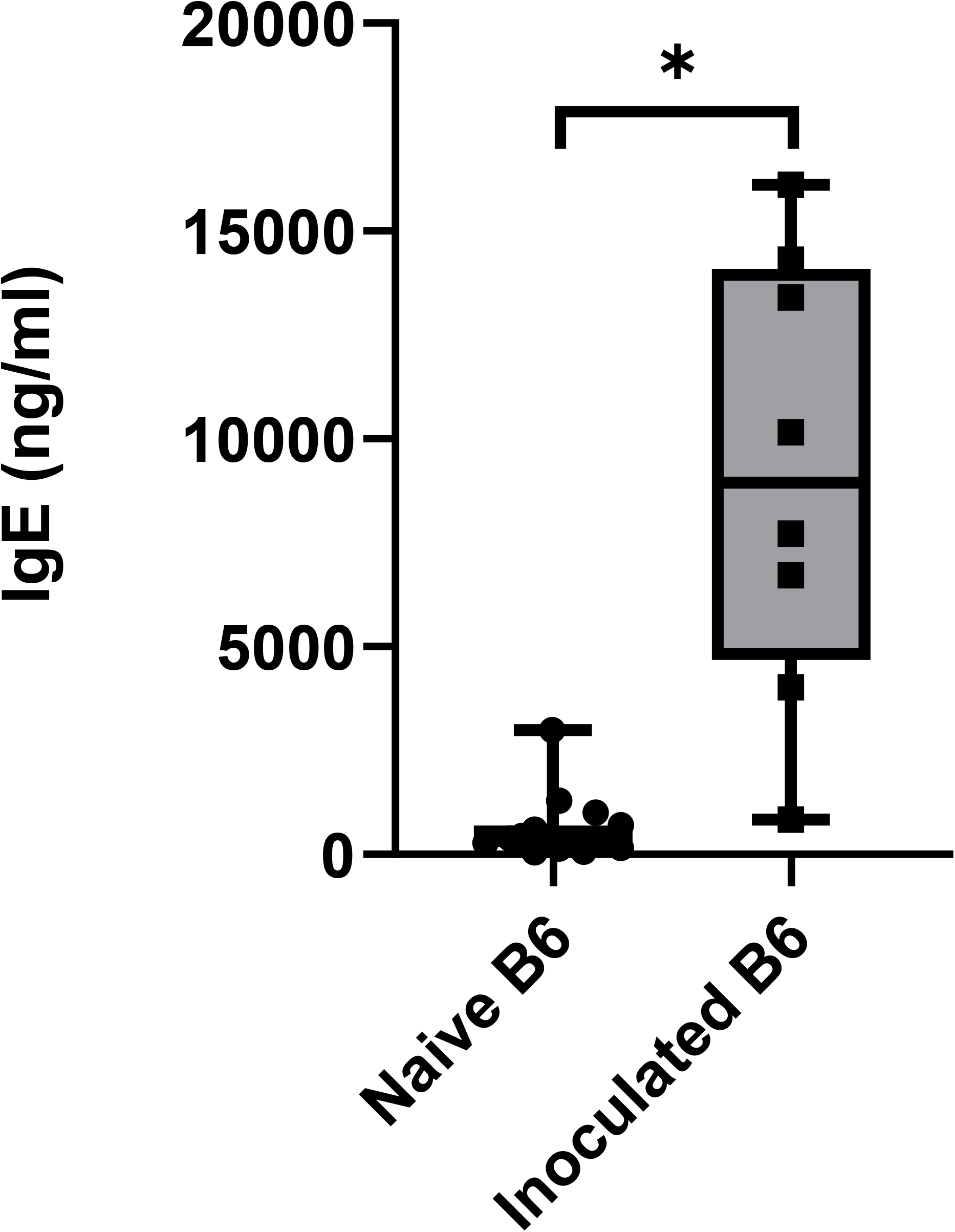
Data shown are the serum IgE concentrations (median [intrabox bar], interquartile range [boxes], minimum and maximum [bars], and individual data points [circles and squares] for naïve B6 mice (n=16; circles) cohoused with Nb- inoculated B6 or NSG mice at 28 days post-housing and Nb-inoculated B6 mice (n= 8; squares) at 35 days post-infection. *IgE values differed significantly (*P* <0.0001) in Nb-inoculated B6 mice compared to naïve B6 mice.

#### Histology

Complete necropsies were performed on Nb-inoculated (B6, n=11; NSG, n=12) and naïve (B6, n=24; NSG, n=24) mice at day 35 PI. On gross necropsy, the lungs from the majority (20 of 23; 87%) of Nb-inoculated mice failed to collapse, were mottled, and had variable degrees of pulmonary emphysema characterized by multifocal gas-filled cystic-like structures on the pulmonary surface. Histopathology was not available for a single Nb-inoculated B6 mouse due to tissue damage during storage. Microscopically, the lungs in 10 of 11 (90.9 %) and 12 of 12 (100%) Nb-inoculated B6 and NSG mice, respectively, showed multifocal to coalescing regions of pulmonary emphysema characterized by alveolar septal wall loss and dilatation (Figure 5 and C). Alveolar histiocytosis, hemorrhage, and hemosiderin-laden macrophages accompanied the lesions in most Nb-inoculated B6 (10 of 11; 90.9%) and NSG (12 of 12; 100%) mice (Figure 5F). Although Nb larvae were rarely observed in the airways of both mouse strains, these lesions are consistent with alveolar damage induced by migrating nematode larvae. Additionally, the lungs of Nb-inoculated B6 mice (8 of 12; 66.7%) often exhibited focal to multifocal areas of perivascular and peribronchiolar inflammation (Figure 5C), whereas this lesion was occasionally observed in the lungs of Nb-inoculated NSG mice (2 of 12; 16.7%%). Terminal bronchioles in a subset of Nb-inoculated NSG cases (6 of 12; 50%) had individual or small clusters of epithelioid macrophages and pyknotic cells containing intracytoplasmic coarsely granular dark-brown pigment (Figure 5E), which reflects prior bronchiolar epithelial damage and hemorrhage after L_3_ migration in the terminal bronchioles. In contrast, none of the naïve B6 and NSG mice cohoused with Nb-inoculated mice had gross abnormalities, alveolar emphysema, pulmonary hemorrhage, or hemosiderin-laden macrophages at sacrifice on 28 DPH (Figure 5). Alveolar histiocytosis was seen in a small number of naïve B6 (4 of 24; 16.7 %) and NSG (8 of 24; 33.3%) mice, and perivascular/peribronchiolar inflammation was identified in 7 of 24 (29.1%) naïve B6 mice.

**Figure 5.**
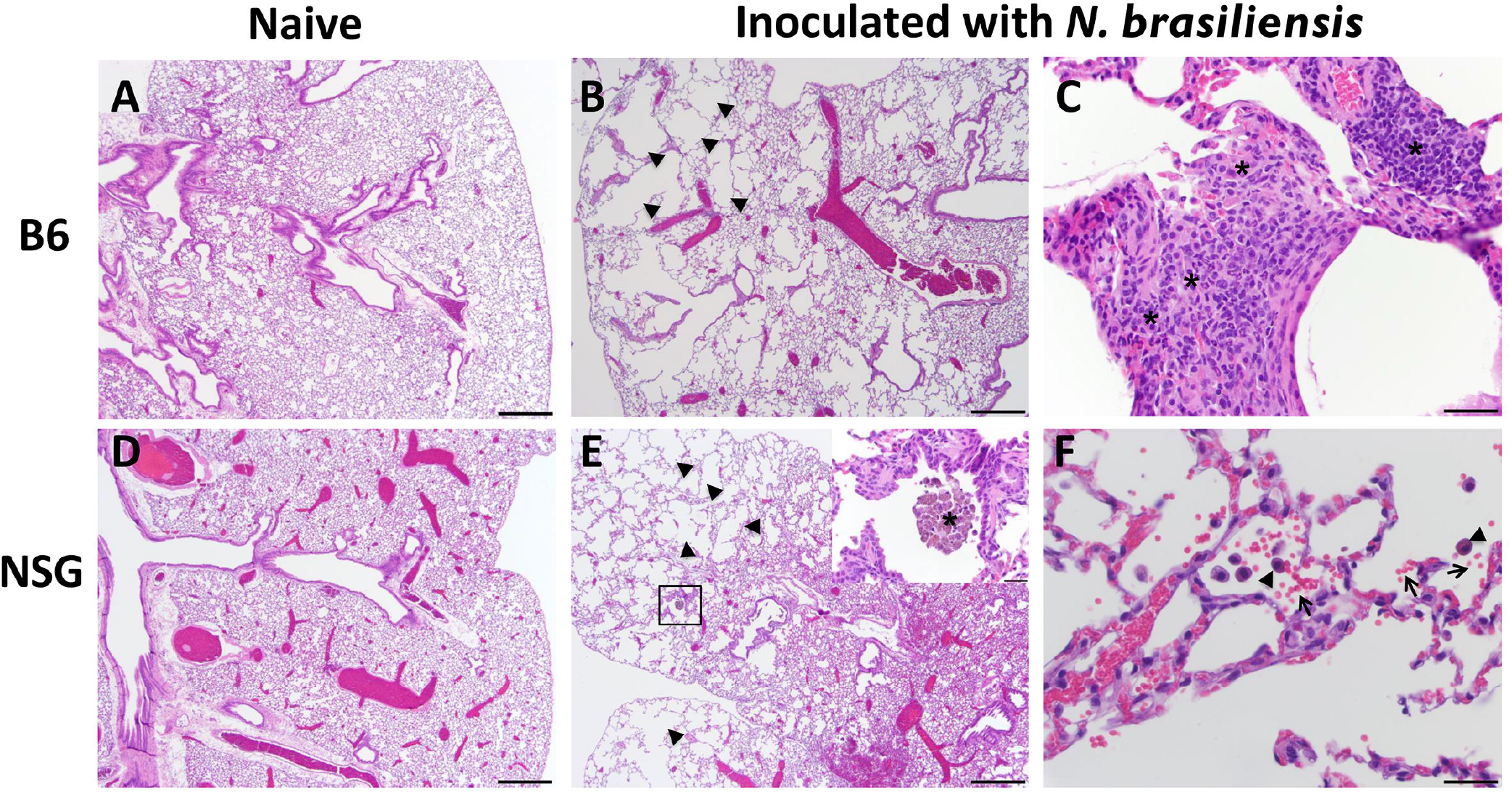
Pulmonary histology of naïve and *Nipostrongylus brasiliensis* (Nb)-inoculated B6 and NSG mice. (A) Normal bronchi/bronchioles and alveoli from a naïve B6 mouse (magnification 4×). (B) Lung from a Nb-inoculated B6 mouse demonstrated marked multifocal to coalescing regions of alveolar emphysema (arrowheads, magnification 4×) with (C) foci of perivascular and interstitial inflammation composed of mixed histiocytic, lymphocytic and neutrophilic infiltrates (asterisk, magnification 40×). (D) Normal bronchi/bronchioles and alveoli from a naïve NSG mouse (magnification 4×). (E) Lung from a Nb-inoculated NSG mouse with marked multifocal areas of alveolar emphysema (arrowheads, magnification 4×) with clusters of epithelioid macrophages and pyknotic cells containing intracytoplasmic coarsely granular dark-brown pigment (asterisks) within the bronchiolar lumens (inset; magnification 40×). (F) The emphysematous alveolar spaces often contained individual hemosiderin-laden macrophages (arrowhead) associated with extravasated erythrocytes (arrows, magnification 40×).

The small intestinal lumens in less than half of the Nb-inoculated NSG mice (5 of 12; 41.7%) contained individual adult nematodes consistent with Nb histomorphology (Figure 6C), while no adult nematodes were noted microscopically in the small intestine of Nb-inoculated B6 mice. Nb eggs were occasionally observed under brightfield microscopy in the cecal and/or colonic content of (3 of 12; 25%) Nb-inoculated NSG mice euthanized day 35 PI (Figure 6F). The small intestine, cecum, colon, heart, lymph nodes, and spleen were histologically normal in all Nb-inoculated and naïve mice.

**Figure 6.**
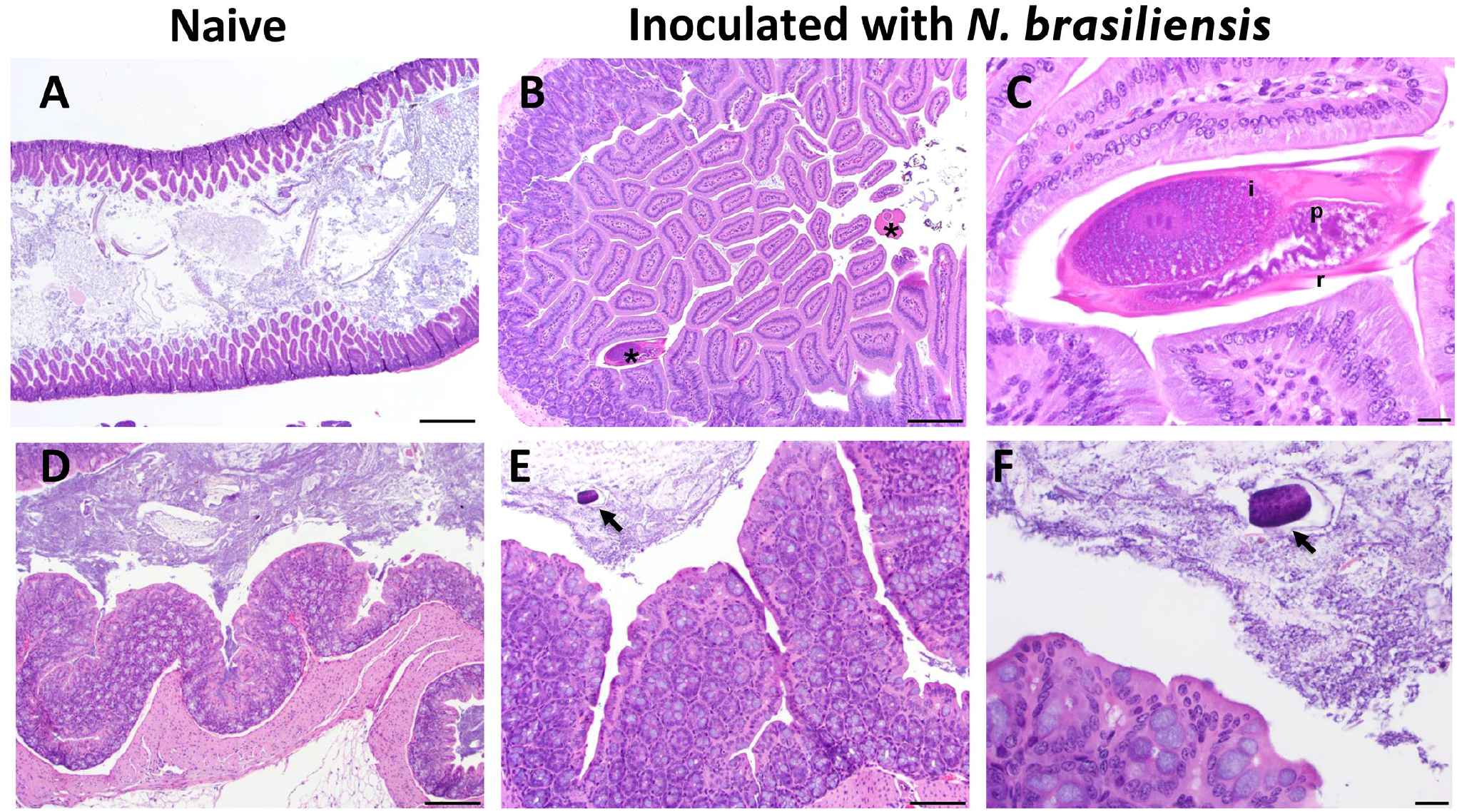
Histology of the small and large intestine of naïve and *Nipostrongylus brasiliensis* (Nb)-inoculated NSG mice. (A) Small intestine from a naïve NSG mouse (magnification 4×). (B) Small intestine from a Nb-inoculated NSG mouse demonstrating 2 Nb adult nematodes (asterisks) associated with normal villous epithelium at day 35 post-inoculation (magnification 10×). (C) Image of a Nb adult (L_5_) nematode with unevenly spaced external cuticular ridges (c) and digestive tract (i) in the pseudocoelom (p, magnification 60×). (D) Colon from a naïve NSG mouse (magnification 10×). (E) Colon from a Nb-inoculated NSG mouse showing a morulated nematode egg (arrow) in the intestinal lumen (magnification 20×). (F) High power image of prior section with nematode egg (arrow) (magnification 60×).

#### Intracage Temperature and Humidity Levels

Temperature and humidity measured in 1 cage from each of 3 randomly selected groups during different cohort repetitions during the 28 days of cohousing were 22.8 +/-0.02°C (mean +/-SEM) and relative humidity of 83.9 +/-0.22%; (mean +/-SEM).

#### Cage Inoculation

No Nb eggs were detected in fecal samples collected for 21 consecutive days from any of the 9 NSG mice housed in cages spiked twice with 10,000 infective Nb L_3_. Neither adult worms nor lesions suggestive of Nb infection were identified at necropsy or on histopathology at 35 DPH.

#### In Vitro Larval Development

Nb eggs developed into infective L_3_ larva under 2 different environmental conditions (moist and control). Mean Nb L_3_ larval counts were higher in moist bedding plates (1095.3 ± 205.3; mean +/-SEM) compared to control plates (709.7 ± 136.5; mean +/-SEM), however the differences were not statistically significant. No L_3_ larvae were found under dry or soiled environmental conditions.

### Supplemental Studies Conducted to Investigate Unexpected Results

#### Nb Inoculation of Athymic Nude Stocks

Egg shedding of the Nb-inoculated nude mouse stocks are shown in Figure 7. Eggs were only detected in 2 of 6 (33.3%) of both nude stocks. Eggs were first detected on 7 DPI in the aforementioned mice. Peak egg shedding occurred on 7 and 8 DPI in nude-1 and nude-2 mice, respectively. Nude-1 mice ceased to shed after 9 DPI, whereas nude-2 mice ceased shedding on 10 DPI. Although the peak number of eggs shed was higher in the nude-1 stock, the cumulative mean EPG was not significantly different between stocks. Grossly, the lungs from the majority (11 of 12; 91.7%) of Nb-inoculated nude mice failed to collapse, were mottled, and had variable degrees of pulmonary emphysema characterized by multifocal gas-filled cystic-like structures on the pulmonary surface. Microscopically, the lungs of Nb-inoculated nude mice (11 of 12; 91.7%) had lesions consistent with Nb larva migration characterized by multifocal to coalescing regions of pulmonary emphysema with alveolar histiocytosis with or without perivascular and peribronchiolar inflammation, hemosiderin-laden macrophages, and hemorrhage. No lesions were noted in the small intestines, cecum, colon, heart, lymph nodes, or spleen on histopathology and no adult nematodes nor eggs were detected in the intestines at necropsy on 35 DPI.

**Figure 7.**
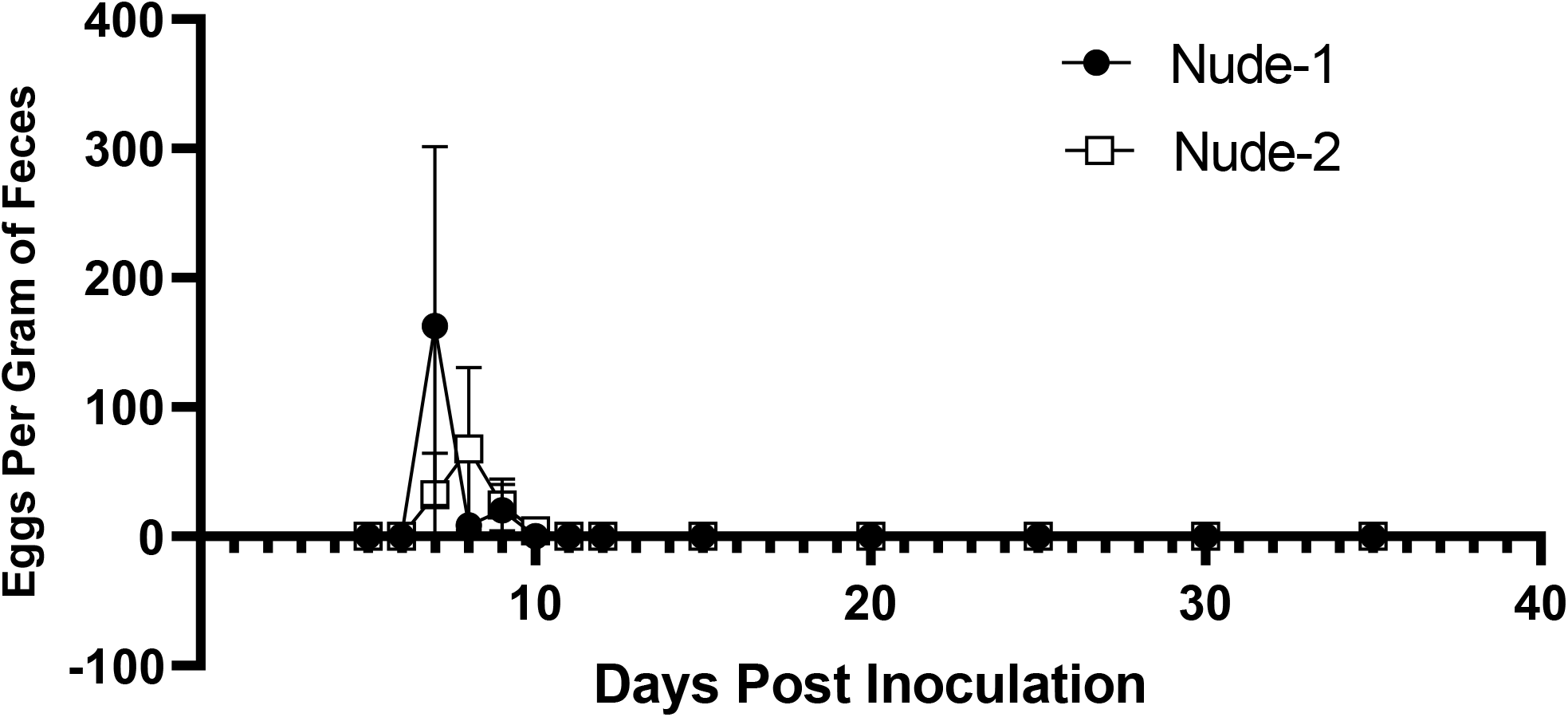
Data shown are the mean (+/-SEM) egg per gram of feces for nude-1 (n=6) and nude-2 (n=6) mice inoculated with Nb L_3_ larvae on day 0 and egg shedding assessed on various days until 35 days post inoculation.

#### Gavage of Feces Containing Nb Eggs

To determine if naïve mice that were unexpectedly shedding eggs had a patent Nb infection, or the eggs detected reflected passage of eggs ingested during coprophagy, 3 NSG mice were gavaged twice with a sugar solution containing approximately 200 Nb eggs. The initial pooled fecal sample was negative. Six eggs were observed in the pooled fecal sample from the NSG mice approximately 16 hours following the second gavage. No additional eggs were detected in feces collected for the next 10 consecutive days, nor were adult worms observed at necropsy or lesions consistent with Nb infection identified on histopathology.

## DISCUSSION

Results from this study demonstrate that horizontal transmission of Nb did not occur, even when highly susceptible mice were cohoused with Nb-infected immunocompromised mice chronically shedding very high numbers of eggs, under optimal conditions for Nb transmission, i.e., ideal intracage temperature and humidity levels in static isolator cages changed at 14-day intervals. Furthermore, highly susceptible NSG mice failed to become infected when housed in cages spiked with large numbers of infective L_3_ larvae and, while eggs were able to develop into infective L_3_ larva under ideal artificial environmental conditions (moist and control), they were unable to develop under conditions created to replicate the intracage environment (dry and soiled).

While eggs were detected in the feces of naïve mice cohoused with Nb-inoculated, egg-shedding NSG mice, we provide data supporting our conclusion that these eggs were presumed to have been ingested during coprophagy as they were detected too early to reflect a patent infection, and mice gavaged with Nb eggs shed eggs without becoming infected. Eggs were only detected in naïve NSG and B6 mice when cohoused with Nb-inoculated NSG mice, which shed large numbers of eggs for the entire period of cohousing. No eggs were identified in naïve NSG and B6 mice when they were cohoused with Nb-inoculated B6 mice, which shed considerably fewer eggs for a short period. When eggs were detected in naïve mice, the quantity shed was limited and shedding was sporadic, and even though eggs were detected in the feces, no adult worms nor lesions consistent with Nb larval migration infection were found at necropsy. In contrast, all Nb-inoculated NSG mice were still shedding eggs and L_5_ nematodes were identified in all mice, and almost all Nb-inoculated B6 mice had histologic evidence of larval migration in their lungs at 35 DPI. Our observations of egg shedding in naïve B6 mice cohoused with Nb-inoculated NSG mice in this study were contrary to what is described in the literature and what we observed in the Nb-inoculated B6 mice; B6 mice typically shed high numbers of eggs for a few days and then cease shedding by 10 DPI.^6; 34; 43^ Additionally, once they are infected and expel the parasite, lifelong immunity is achieved.^5; 6; 19^ During our study, eggs were initially detected in the feces of naive B6 mice on day 7 DPH and were detected intermittently until 28 DPH. Nb eggs take approximately 5 to 6 days to embryonate and hatch.^5; 6; 18; 25^ As the detection of eggs on 7 DPH was surprising, fecal egg counts were initiated at 1 DPH in subsequent cohorts, a time point in which eggs were still detected, providing further evidence that the eggs were passing through the gastrointestinal tract following ingestion of egg-containing feces during coprophagy. Rats experimentally infected with low doses of L_3_ larvae daily over weeks, to mimic natural exposure, shed eggs persistently but they also had L_5_ Nb in their intestines at necropsy.^23; 24^ To confirm that our hypothesis was correct, NSG mice were gavaged with eggs collected from fresh fecal pellets obtained from Nb-inoculated NSG mice. A small number of eggs were identified in fecal pellets collected from mice the day following gavage. No additional eggs were found in fecal samples collected for 10 consecutive days, and no adult worms nor were microscopic lesions suggestive of Nb infection found at necropsy. Interestingly, the mice needed to be gavaged twice, on consecutive days, as eggs were not detected following the first administration. This result was surprising, and we can only speculate as to the reasons, which include the potential that the eggs degraded in the gastrointestinal tract and/or the small sample size evaluated may have resulted in an initial false negative result.

Nb infection is known to induce production of polyclonal and specific IgE.^5; 6; 32; 42^ Elevations in total serum IgE levels in Nb-inoculated BALB/c mice are seen within 3 to 5 days following infection, peak at 10 to 17 days, and remain elevated for over 60 days.^42^ One study demonstrated that total IgE serum concentrations in Nb-inoculated BALB/c mice reached a maximum of 200 µg/ml at 15 DPI, but levels declined to 40 µg/ml by day 25 DPI.^27^ In B6 mice, total serum IgE levels measured 14 days following inoculation were 2,060 ± 753 (mean +/-SD).^28^ In our study, total IgE concentrations in Nb-inoculated B6 mice differed from what has been previously reported; however, mouse strain, collection time points, and possibly assay differences likely account for the variations. It is well known that the B6 strain mounts a Th1 dominant response in contrast to the BALB/c strain whose response is Th2-skewed.^35^ Rats administered low doses of Nb L_3_ larvae, meant to simulate natural infection, develop significant and sustained IgE responses.^41^ Therefore if horizontal transmission occurred, we would have expected to detect markedly elevated IgE levels in naïve B6 mice. While an elevated serum IgE concentration was detected in a single naïve B6 mouse, no eggs were detected during 28 days of cohousing, neither lesions consistent with Nb infection nor were nematodes detected at necropsy in this mouse. IgE concentrations may be elevated for a variety of reasons, including biological variation and exposure to allergens. In addition, the finding that IgE levels in B6 naïve mice with Nb eggs in their feces were significantly lower than those in Nb-inoculated mice, provides further support that these eggs were ingested and not a reflection of patent infections.

While Nb infections have been studied in immunocompromised mouse strains, this study, to our knowledge, reflects the first time that the infection has been characterized in the severely immunocompromised NSG strain.^1; 4; 10; 19; 21; 22; 31; 34; 36^ NSG mice reached peak egg shedding at 9 DPI, and shed markedly more eggs as compared to B6 mice during the study period. While the quantity of eggs shed did decrease over time, NSG mice continued to shed high numbers of eggs throughout the study suggesting that they would likely remain persistently infected.

Athymic nude mice were originally selected for this study to serve as the immunocompromised strain due to their reported ability to shed high numbers of eggs for a protracted period as they are chronically infected.^1; 14; 21; 22; 31^ However, eggs were only detected in 2 of 6 nude-1 mice, the number of eggs shed was low, and shedding was sporadic with cessation at 9 DPI. To determine if these findings were a result of genetic differences, a different nude stock from an alternate vendor (nude-2) were inoculated with the same number of L_3_ larvae yielding similar results. Pulmonary lesions, consistent with larval migration, were identified in 91.7% of the athymic nude mice with no discernable differences in lesion severity observed between the stocks. While patent infection did occur, no L_5_ Nb were detected at 35 DPI. While resistance to Nb infection has been reported in FVB/N mice and IL-5 transgenic mice, it has not been described in athymic nude mice.^11; 12; 26^ Pulmonary lesions consistent with Nb infection and low to absent egg counts suggest damage to larvae likely occurred during early stages of infection implicating the innate immune system as a potential source of resistance.^11; 12; 26^ Athymic nude mice have been shown to be resistant to some pathogens including experimental cutaneous infection with *Bacillus anthracis, Candida albicans, Listeria monocytogenes*, and *Brucella abortus*.^7; 13; 17; 39^ Proposed mechanisms for resistance include potent neutrophilic responses, augmented effects of prior microbial exposure in stimulating the innate immune system due to lack of T-cell regulatory responses, and enhanced activated macrophage functions.^7; 13; 17; 39^ The differences between our observations and earlier studies using the athymic nude mouse could be attributed to the use of inbred versus outbred nude mice, the number of larvae administered, and the route of inoculation.^1; 14; 21; 22; 31^ Parasite infectivity and worm burden can also be affected by the strain of nematode used.^5^ The number of passages and time since hatching can also contribute to changes in parasite viability and variations in worm burden.^5^ Serial passage of Nb through rats rather than mice could also have impacted infection kinetics in athymic nude mice.^40^

The pulmonary changes seen in all Nb-inoculated mice were consistent with the chronic phase of Nb infection, which is characterized by pulmonary emphysema with alveolar hemorrhage and hemosiderin-laden macrophages in the absence of an overt inflammatory leukocytic cell infiltrate.^5; 29; 33^ Damage to lung tissue occurs as the L_3_ larvae migrate from blood vessels into alveolar spaces during early stages of Nb infection.^5; 8; 9; 29^ Mechanical tissue destruction during larval migration causes leakage of erythrocytes into alveolar spaces, which is resolved by the recruitment of alveolar macrophages and erythrophagocytosis.^5; 8^ In the mouse model, Nb-induced emphysema is typically studied at around 30 DPI.^5^ For our study we sought to compare histopathologic lesions in Nb-inoculated mice with cohoused naïve mice in order to determine if horizontal transmission occurred. No lesions consistent with chronic Nb infection were identified in the lungs of any naïve mice. Alveolar histiocytosis was identified in a small number of naïve B6 and NSG mice and perivascular and peribronchiolar inflammation was identified in about a third of the naïve B6 mice. These lesions are incidental background lesions described in these strains and are not suggestive of Nb infection.

Small intestinal inflammation, villous atrophy, flattening and fusion, and crypt hyperplasia have been described in heavy Nb infections.^3; 25^ However, small intestinal lesions were not noted in Nb infected or naïve mice. In this study the proximal half of the small intestine was removed to perform nematode counts and thus was not examined microscopically as L_5_ Nb typically inhabit the proximal portion of the small intestine. That may explain why small intestinal lesions were not observed and why only a few adult worms were found in the small intestines of Nb infected NSG mice on histopathology. Another reason lesions may have been absent in the gastrointestinal tract is that necropsies were performed in Nb-inoculated B6 mice at 35 DPI as opposed to the period of active infection 6 to 10 DPI.^5; 6; 34; 43^

NSG mice were housed in cages spiked with 10,000 infective L_3_ larvae on days 7 and 21 after housing to model a “worst-case” scenario with a high likelihood that larvae would penetrate the skin resulting in infection. Larvae were initially added to the cage 7 days following housing to ensure that the bedding was sufficiently moist to support larval survival promoting skin penetration. Fecal flotation performed daily after the cages were initially spiked remained egg-free. Additionally, no adult L_5_ Nb or lesions suggestive of infection were identified at necropsy.

We demonstrated that while eggs were able to develop and hatch releasing L_3_ larvae under extremely moist environmental conditions, larvae were incapable of developing under environmental conditions likely to be found within mouse cages while providing the optimal temperature and humidity for Nb development.^5; 6; 25^ Surprisingly, more L_3_ developed in moist autoclaved aspen chip bedding as compared to conditions used to propagate larvae in the laboratory. These differences may have resulted from variation in the number of eggs in individual fecal pellets used to inoculate the plates. Findings indicated that moisture plays a crucial role in the ability of eggs to hatch and develop into L_3_ larvae. While soiled plates did contain moisture, urine has an adverse effect on eggs which may explain why L_3_ larvae did not develop under these conditions.^6^

One unexpected challenge faced in this study was the inability to use fecal flotation to confirm horizontal transmission. Egg counts are typically used to confirm and quantify infection in Nb-inoculated mice, however in this study it was ineffective due to coprophagy.^6^ Another challenge was that as mice were cohoused for 28 days to allow sufficient time for horizontal transmission to occur, there was no definitive day of infection on which time points for larval detection in the skin or lungs or nematodes in the small intestine could be focused.^2; 5; 6; 18; 25^ One could consider a limitation of this study was the fact that immune response evaluation was limited to total serum IgE in B6 mice. IgE was measured as it remains elevated for an extended period following infection.^15; 27; 42^ While IgE was not measured in NSG mice, IL-5 levels could have been evaluated. Natural and experimental infection with Nb induces a profound CD4^+^ Th2 dependent immune response characterized by production of IL-4, IgE, IL-13, type 2 innate lymphoid cells (ILC2s), Th2 thymocyte expansion, IL-5 dependent eosinophil recruitment and alternatively, activated macrophages (AAMs), IL-10 production, as well as goblet cell hyperplasia and mastocytosis.^5; 6; 37; 38^ Further analysis of the lungs and gastrointestinal tract could have been performed to examine changes in T-cell, eosinophil, innate lLC2, and cytokine populations.^15; 30^ We also used a rat adapted Nb strain which could have affected infectivity kinetics, however rat adapted Nb are used by many laboratories using the parasite to interrogate the mouse’ s immune system, and we inoculated mice with the recommended and commonly used higher larval dose.^6; 40^

In summary, this study provides compelling objective evidence that horizontal transmission of Nb does not occur under the various scenarios that were modeled to provide optimal conditions for its transmission. These results led our institution to revisit biosecurity practices, which had historically subjected murine models of Nb infection to be conducted at ABSL-2, to currently employ ABSL-1 practices. Findings from this study can be used by others to inform the decision as to the appropriate biosecurity measures to be implemented when using the Nb mouse model at their institution.

## Abbreviations and Acronyms

Nb: Nippostrongylus brasiliensis
L_3_: Infective larvae (3^rd^ stage)
L_5_: Adult worms
DPI: Days post inoculation
DPH: Days post cohousing
EPG: Eggs per gram of feces
AAM: Alternatively activated macrophages

## Acknowledgments

We thank Beatrice Hoyos and the Rudensky Laboratory for their donation of *Nippostrongylus brasiliensis* L_3_ larvae and for assistance in larval culture, Jacqueline Candelier and the staff of the Laboratory of Comparative Pathology for histology support, Juliette Wipf for running IgE assays, Sara Mangosing for assistance with larval preparation and inoculation, and Enrico Capobianco, Ji-Gang Zhang, and Andrew Schile for assistance with statistics. MSK Core Facilities are supported by the NCI Cancer Center Support Grant P30 CA008748.

